# Bridging microscopic and macroscopic mechanisms of p53-MDM2 binding using molecular simulations and kinetic network models

**DOI:** 10.1101/086272

**Authors:** Guangfeng Zhou, George A. Pantelopulos, Sudipto Mukherjee, Vincent A. Voelz

## Abstract

Under normal cellular conditions, the tumor suppressor protein p53 is kept at a low levels in part due to ubiquitination by MDM2, a process initiated by binding of MDM2 to the intrinsically disordered transactivation domain (TAD) of p53. Although many experimental and simulation studies suggest that disordered domains such as p53 TAD bind their targets nonspecifically before folding to a tightly-associated conformation, the molecular details are unclear. Toward a detailed prediction of binding mechanism, pathways and rates, we have performed large-scale unbiased all-atom simulations of p53-MDM2 binding. Markov State Models (MSMs) constructed from the trajectory data predict p53 TAD peptide binding pathways and on-rates in good agreement with experiment. The MSM reveals that two key bound intermediates, each with a non-native arrangement of hydrophobic residues in the MDM2 binding cleft, control the overall on-rate. Using microscopic rate information from the MSM, we parameterize a simple four-state kinetic model to (1) determine that induced-fit pathways dominate the binding flux over a large range of concentrations, and (2) predict how modulation of residual p53 helicity affects binding, in good agreement with experiment. These results suggest new ways in which microscopic models of bound-state ensembles can be used to understand biological function on a macroscopic scale.

**AUTHOR SUMMARY:** Many cell signaling pathways involve protein-protein interactions in which an intrinsically disordered peptide folds upon binding its target. Determining the molecular mechanisms that control these binding rates is important for understanding how such systems are regulated. In this paper, we show how extensive all-atom simulations combined with kinetic network models provide a detailed mechanistic understanding of how tumor suppressor protein p53 binds to MDM2, an important target of new cancer therapeutics. A simple four-state model parameterized from the simulations shows a binding-then-folding mechanism, and recapitulates experiments in which residual helicity boosts binding. This work goes beyond previous simulations of small-molecule binding, to achieve pathways and binding rates for a large peptide, in good agreement with experiment.

## Introduction

The transcription activator p53 plays a central role in tumor suppression^1^. Cellular levels of p53 are normally kept low by targeted degradation by the E3 ubiquitin ligase MDM2 (mouse double minute 2), whose N-terminal domain binds residues 17-29 of the p53 transactivation domain (TAD) in a deep hydrophobic cleft^2^. The p53 TAD is intrinsically disordered^3^, but forms a helix when bound to MDM2. Various types of cellular stresses such as DNA damage leads to disruption of p53-MDM2 binding and an increase in p53 expression, which in turn promotes cellular repair or apoptosis. Thus, the discovery of potent competitive inhibitors that can disrupt the p53-MDM2 binding interaction has been an important strategy for developing new cancer therapeutics^4,5^. The availability of structural information has also made p53-MDM2 a valuable model system for the study of protein-protein interactions and the development of new classes of peptidomimetics^6–9^ often alongside computational design efforts^10,11^.

A consensus of experimental and simulation studies suggest that intrinsically disordered protein (IDP) domains such as p53 TAD bind their receptors through an induced-fit “fly-casting” mechanism, whereby binding occurs first, followed by structuring into higher-affinity poses^12–17^. It has been proposed that this mechanism facilitates binding to multiple partners in complex regulatory networks, and may enable fast association rates important for signaling. The structural and kinetic properties of IDPs are thought to fine-tune many signaling interactions^18^. Recently, Borcherds et al. have shown that the extent of residual helicity of p53 TAD can modulate p53-MDM2 binding affinity as well as signaling dynamics in cells^19^. An important challenge for molecular simulation is thus to predict binding pathways and association rates of IDPs to their targets, and the detailed molecular mechanisms responsible for shaping them.

In this work, we use extensive all-atom molecular simulations in explicit solvent, combined with state-of-the-art Markov State Model (MSM) approaches, to investigate the p53-MDM2 binding mechanism. Recent MSM studies have examined the mechanisms by which protein receptors recognize small molecules^20–23^ and here we extend similar methods to model the coupled folding and binding of a larger peptide (p53 TAD peptide) to MDM2.

## Methods

### Molecular simulation

Simulations starting from unbound states were performed on the the Folding@home distributed computing platform^24^ with Gromacs 4.5.4^25^ using the Amber ff99sb-ildn-nmr force field^26^ and TIP3P explicit solvent. A number of initial starting configurations were selected from conformational clustering of implicit-solvent REMD simulations, each placed at different distances within 12 Å from the binding site. A total 2776 trajectories were generated amounting to ~ 831*μs* of aggregate simulation data (Figure S1).

### MSM construction

Recent methodological advances have exploited the variational approach to conformational dynamics^27^ to enable the construction of optimal MSMs given the available trajectory data^28,29^. To implement this approach, we used time-structure-based Independent Component Analysis (tICA)^30,31^ to project the trajectory data to a low-dimensional subspace that best preserves the slowest conformational transitions. Using all pair distances between *C*_*α*_ + *C*_*β*_ atoms of p53 and the binding pocket of MDM2 (see Supporting Information for details), we constructed a time-lagged correlation matrix **C**^(Δ*t*)^ and corresponding covariance matrix from the pair distances using a Δ*t* = 5 ns lagtime. The tICA components *α* are found by maximizing the objective function ⟨*α*_*i*_|**C**^(Δ*t*)^|*α*_*i*_⟩ subjected to certain constraints^30^. Once projected to the tICA subspace, distance-based clustering using the *k*-means algorithm was performed to obtain MSM metastable state definitions. To select hyper-parameters such as the number of tICA components, clustering method, MSM lag time, and the number of MSM microstates, we performed variational crossvalidation using the GMRQ method of McGibbon et al.^28^ on over 120 MSMs. As in previous work^32^, we find that tICA distance metrics are better than rmsd or dihedral angle metrics, and *k*-means clustering performs better than *k*-centers. The optimal MSM, used in all subsequent analysis, is constructed using 10 tICA components, 600 microstates and a 5 ns MSM lag time τ_lag_ (Figure S2). The MSM transition matrix **T**(τ_lag_) was estimated using a maximum likelihood method^33,34^. The model is validated by implied timescales which plateau near the chosen lag time of 5 ns (Figure S3), and Chapman-Kolmogorov tests (Figure S4). All models were built using MSMbuilder 3.3^33^ and MDTraj 1.5^35^ software packages.

## Results

*Binding precedes folding*. A projection of the simulation data to two reaction coordinates-the rmsd of p53 to its native structure, and the distance of p53 to the MDM2 binding pocket-suggests that binding of p53 precedes folding of p53, consistent with the “fly-casting” mechanism (Figure 1). The distance vs. rmsd landscape can be manually partitioned into four states: folded-bound (blue), unfolded-bound (green), folded-unbound (red), and unfolded-unbound (cyan). These states were defined using a bound-state distance cutoff of 1.2 nm, and rmsd cutoff of 0.2 nm. Projecting the 600 MSM microstates to this landscape, we find most of the population in the bound states, with only one microstate corresponding to the folded-unbound state (red).

**FIG. 1.**
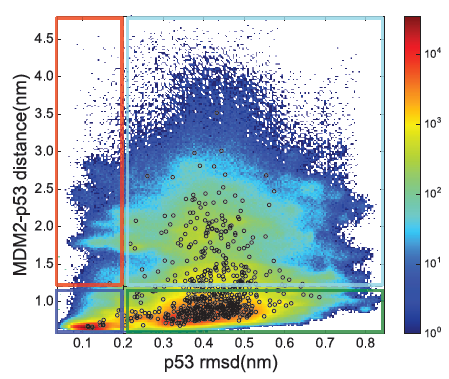
Simulation trajectory data projected to p53-MDM2 distance and the p53 rmsd-to-native show a preference for binding before folding. Colored boxes highlight the partitioning into four states based on these reaction coordinates: folded-bound (blue), unfolded-bound (green), folded-unbound (red), and unfolded-unbound (cyan). Black circles show the locations of cluster centers of the 600 MSM microstates.

Projections of the simulation data to the two largest tICA components, corresponding to the slowest conformational dynamics, show a very different landscape (Figure 2). The folded-bound state (blue) is composed of a single well-populated basin, closely matching within 1.3 Å backbone rmsd to the native co-crystal structure, with the side chains of F19, W23 and L26 correctly inserted into the binding pocket of MDM2. In contrast, the unfolded-bound state (green) is distributed throughout the tICA landscape. The two predominant basins of the unfolded-bound state correspond to p53 bound in two different misfolded states, each with residue F19 in its native binding groove, but W23 outside of the binding cleft. As can be seen by the eigenvector structure of the MSM (Figure S5) transitions from these basins control the slowest timescales of binding.

**TABLE I.** Comparison of experimental and simulated rates.

| Methods | $k_{\text{on}}$ ($\text{M}^{-1} \text{s}^{-1}$) | $k_{\text{off}}$ ($\text{s}^{-1}$) |
| --- | --- | --- |
| Experiment<sup>39</sup> | $9.2 \times 10^6$ | 2.0 |
| TPT 600-state MSM | $2.5 \times 10^7$ | $1.9 \times 10^5$ |
| TPT 4-state MSM | $5.2 \times 10^7$ | $6.0 \times 10^5$ |

**FIG. 2.**
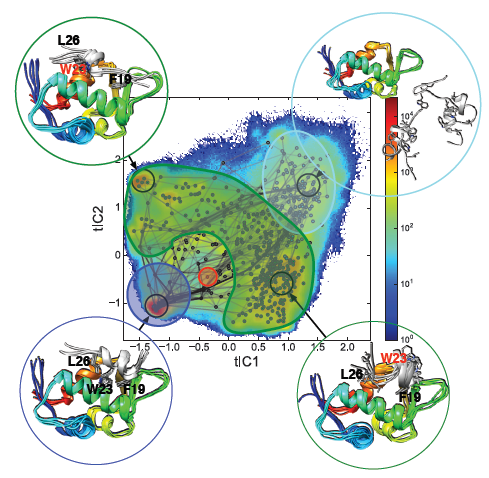
Trajectory data projected to the 2D tICA landscape, with microstates color-coded and overlaid with color-shaded cartoons to highlight the locations of the four states. The 100 highest-flux binding pathways calculated from TPT (black lines) fall into two groups, each dominated by a different unfolded-bound basin with W23 misregistered outside of the binding cleft. Representative structures for these two basins, the folded-bound native state, and the unfolded-unbound state are shown in circles.

*Transition pathways and rates*. To estimate pathways, fluxes and rates of p53 association, we used Transition Path Theory (TPT), which we briefly describe here and refer readers to other references for more details^36–38^. In TPT, source states (*A*) and sink states (*B*) first need to be defined for the transition process of interest. The remaining states are considered to be intermediate states (*I*). Next, committor probabilities 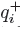, defined as the probability that a trajectory started from state *i* will reach *B* before state *A*, are computed from the MSM transition matrix. The total folding flux giving the expected number of observed *A* → *B* transitions per time unit τ is: 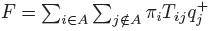. The rate of reaction *A* → *B, k_AB_* can then be computed as:

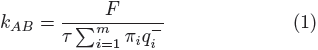

We first used TPT to estimate overall p53 binding on-and off-rates (*k*_on_ and *k*_off_) from the 600-microstate MSM, using the unfolded-unbound and folded-bound states as the source and sink states for *k*_on_, respectively (and vice versa for estimating *k*_off_). For comparison, we constructed a four-macrostate MSM by manually lumping the 600 microstates according to our four-state definitions. The results (Table I) show that the *k*_on_ estimated from the 600-microstate MSM is very close to the experimental *k*_on_ (within a factor of 2.7). Macrostate lumping into four states further accelerates the dynamic timescales, with *k*_on_ predictions still within a factor of 5.6. As expected given the available trajectory lengths, *k*_off_ estimates from both models are severely over-estimated.

*A four-state kinetic model predicts an induced-fit mechanism for p53 binding*. To analyze whether the binding mechanism follows a conformational selection (CS) or induced fit (IF) mechanism (or aspects of both), we computed the reactive flux for each mechanism according to the method introduced by Hammes et al.^40^, illustrated in Figure 3. In this model, association of a ligand can occur either through a weak-binding (*w*) form of p53, or a tight-binding (*t*) form, with interconversions between these two possible when unbound or bound. Here, we slightly modify our interpretation of the model for use with disordered peptide binding; in our case it is the *receptor* MDM2 which can select or induce folded (helical) states of p53 TAD peptide. Following Hammes et al., we compute the reactive flux for conformational selection pathways as:

**FIG. 3.**
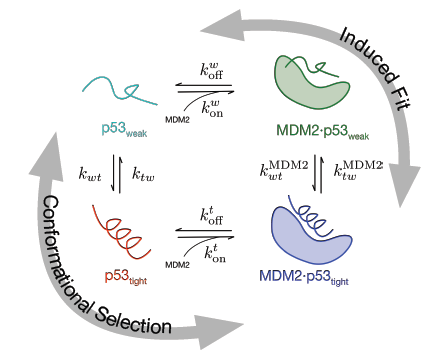
A four-state model of possible p53-MDM2 binding mechanisms includes both induced-fit (IF) and conformational selection (CS) pathways.

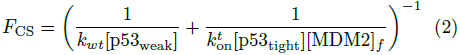

and the flux for induced fit pathways as

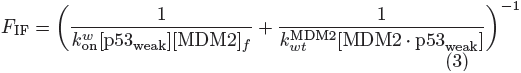

where [MDM2]_*f*_ is the free MDM2 concentration. The derivation is shown in the Supporting Information.

To obtain the relative amounts of reactive flux that occur by conformational selection vs. induced fit pathways, we use our MSM model to make initial estimates of all eight rates in the four-state kinetic model shown in Figure 3. We tried two different approaches to make these initial estimates: (1) directly from the transition probabilities of a four-macrostate MSM derived for our state classifications, and (2) using TPT with pairs of relevant states selected as the source and sink states. Both sets of estimates are listed in Table S1. We find that the two methods yield very similar results, except that 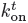 estimates from the transition matrix are more than an order of magnitude larger than the estimate from TPT, due to enforcing detailed balance with a low equilibrium population predicted for the folded-unbound state. Therefore, in the following analysis we use only the parameters estimated from TPT. From this initial estimate, we then scale the off-rates 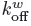 and 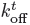 to reproduce the experimental binding affinity of p53 to MDM2 (see Supporting Information).

Unlike our MSM, which was constructed from simulations performed at a fixed concentration ([p53] = [MDM2] = 7.1 mM), the resulting four-state kinetic model can be used to extrapolate the binding fluxes at any desired concentrations. In all cases, we find that binding is dominated by an induced-fit mechanism, consistent with “fly-casting”. The fraction of flux that occurs by an induced-fit mechanism, *F_IF_/(F_CS_ + F_IF_)* is nearly 100% regardless of the concentrations of p53 and MDM2. This is mainly due to the fact that the simulated helicity of p53 is very low (0.11%, Figure 4).

**FIG. 4.**
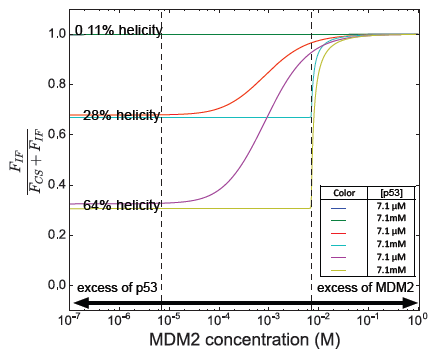
Fraction of binding flux going through induced-fit (IF) pathways at different p53 concentrations and helicities versus total MDM2 concentration. Dashed lines denote the p53 concentrations shown. 7.1 mM is the concentration of p53 and MDM2 in the molecular simulations.

*Increased residual helicity leads to enhanced p53 binding and a shift toward conformational selection*. To estimate the effect of residual helicity of p53 on binding mechanism and affinity, we use a maximum caliber approach to infer how the rates *k_wt_* and *k_tw_* between unfolded-unbound states (cyan) and folded-unbound states (red) change in response to new helix-coil equilibrium populations (see Supporting Information), using the relation 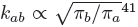. The helicity of p53 predicted by our 4-state model is 0.11%. To model the experimental system of Borcherds et al., we increase the helix population of unbound p53 to 28% and 64%, values measured for wild-type and P27A variants of the p53 TAD^19^. The inferred rates are shown in Table S3.

The four-state kinetic model predicts that increasing residual helicity increases the flux of conformational selection at low MDM2 concentration; however, in the limit of excess of MDM2, induced-fit binding flux increases to almost 100% (Figure 4). The reason for this is the relatively high 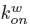 value, a key feature of intrinsically disordered proteins that we have calculated directly from the MSM model. In the limit of excess of MDM2, 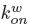 would have to be reduced by several orders of magnitude to convert the binding mechanism to conformational selection. In the limit of excess p53, a shift towards a conformational selection binding mechanism is observed, although a strong preference for induced-fit binding pathways (more than 30% of the binding flux) remains even at high levels of residual helicity (64%) and in the excess of p53.

In agreement with experiment, the four-state model predicts a greater apparent binding affinity of P27A p53 TAD compared to wild-type, with absolute and relative binding free energies similar to experimental values (see Figure 5). The predicted apparent ΔG of binding for p53 wild-type and P27A are −7.5 and −9.0 kcal·mol^−1^, respectively, while the experimental values are −9.1 ± 0.2 and −10.4 ±0.1 kcal·mol^−1^, respectively. We predict that the ΔΔG incurred by increasing the helicity of p53 from 28% to 64% is −1.5 kcal·mol^−1^, which also agrees very well with experiment (−1.3±0.3 kcal·mol^−1^).

**FIG. 5.**
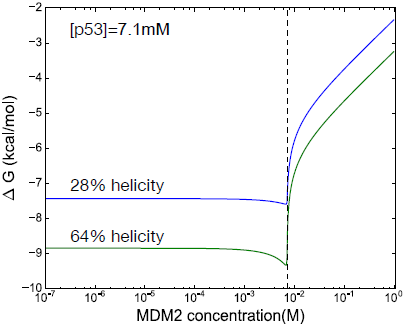
Predicted p53-MDM2 binding affinities versus MDM2 concentration from the four-state kinetic model. Increasing the residual helicity of p53 TAD from 28% to 64% increases affinity by ΔΔ*G* = −1.5 kcal/mol, in good agreement with experiment (−1.3 ± 0.3 kcal/mol).

## Discussion

Recently, Zwier et al. has reported efficient implicit-solvent simulations of p53 TAD peptide binding to MDM2 carried out using a weighted-ensemble path sampling strategy^42^ on 3500 CPU cores of TACC Stampede for 15 days, with an aggregate simulation time of 120 μs.^43^ The authors report similarly accurate predictions of on-rates, and a mechanism whereby diffusion-controlled formation of a specific encounter complex is the rate-limiting step. While their study predicts a high helical propensity for the p53 TAD peptide, our study predicts less helicity, possibly due to differences in the forcefield and solvent model used, as well as differences in initial starting conformations.

An advantage of our approach of parameterizing a four-state binding flux model from a detailed MSM is the ability to extrapolate differences in binding mechanisms that result from various helical propensities of p53 TAD peptide and various ligand and receptor concentrations. At the effective concentrations used in our simulation, residual helicity exerts a large influence on dominant binding flux, but has less influence in excess MDM2 (see Figure 4).

Notably, both our study and the Zwier et al. study suggest a bright future using adaptive sampling simulations to model protein-peptide binding. With a high-quality MSM of p53 binding now constructed, we aim to exploit new MSM-based adaptive sampling approaches to model binding rates and mechanisms for multiple sequences^44^. A remaining challenge of course is to efficiently sample off-rates as well as on-rates. With the advent of multiple-ensemble MSM techniques^45^, this too may be within reach in the near future. We expect that the mechanistic detail provided by MSM approaches may suggest new ways to design inhibitors that compete with natural substrates.

## Conclusion

We have used *ab initio* binding simulations and Markov State Models to construct a detailed kinetic network model of p53 TAD peptide binding to MDM2. The MSM predicts binding on-rates in agreement with experiment, as well an ensemble of encounter complex structures that control the overall binding pathways and rates. Predicted MSM rates, along with experimental affinities, were used to parameterize a four-state kinetic model, which predicts an induced-fit “fly-casting” mechanism over a wide range of concentrations, and shows increased binding affinity for p53 variants with higher amounts of residual structure, in agreement with recent experiments. This work demonstrates how combining detailed all-atom MSMs and simple few-state kinetic models can be very useful in understanding how disordered protein domains bind their target receptors. The results also suggest new ways to design inhibitors that compete with natural substrates, by rationalizing how specific binding modes may modulate key rate processes, in the context of physiological concentrations.

## Acknowledgments

The authors thank the participants of Folding@home, without whom this work would not be possible. We thank Dr. Lillian Chong for very helpful feedback on our manuscript. This research was supported in part by the National Science Foundation through major research instrumentation grant number CNS-09-58854 and MCB-1412508.

## Supporting Information

### Supporting Text

*Preparation of initial p53-MDM2 configurations for simulation*. Four non-native structures of the p53 fragment were chosen from 500 ns of replica exchange molecular dynamics (REMD) simulations of the p53 fragment (residues 17-26) taken from PDB structure 1YCR. The Amber ff96 force field with the OBC GBSA implicit solvent model were used to simulate 16 exponentially-spaced temperature replicas between 300 and 450 K. Conformational clustering was performed on the lowest-temperarture replica using a backbone-RMSD distance metric, from which representative structures of p53 were chosen. Of these structures, two were partial helical, one were hairpin-like and one was random coil. These four p53 TAD peptide structures, in addition to the native structure, were used to prepare 30 initial structures of the p53-MDM2 complex at a variety of distances between the p53 fragment and the MDM2 binding pocket by placing six replicas of each p53 fragment at 2 Å intervals (2, 4, 6, 8, 10 and 12 Å) along a vector from the MDM2 center of mass to the p53 center of mass the 1YCR crystal structure.

*Pairwise atom distances used for time-lagged independent component analysis (tICA)*. To perform tICA, the simulation trajectory data was first projected to a set of 1953 distance coordinates, from which a time-lagged correlation matrix was constructed. We used all pairwise distances between C_*α*_ and C_*β*_ atoms in selected residues of p53 (Glu17, Thr18, Phe19, Ser20, Asp21, Leu22, Trp23, Lys24, Leu25, Leu26, Pro27, Glu28, Asn29) and MDM2 (Glu25, Thr26, Met50, Lys51, Leu54, Leu57, Gly58, Ile61, Met62, Tyr67, Gln71, Gln72, His73, Val75, Phe91, Val93, Lys94, His96, Ile99).

*Distance between p53 and the MDM2 binding pocket*. Figure 1 in the main text uses as a structural observable the distance of p53 to the MDM2 binding pocket. We define this distance as the average of six p53-MDM2 pairwise C_*α*_ distances: (p53-Phe19, MDM2-Met62), (p53-Phe19, MDM2-Gln72), (p53-Trp23, MDM2-Gly58), (p53-Trp23, MDM2-Val93), (p53-Leu26, MDM2-Leu54), and (p53-Leu26, MDM2-His96). Selected residues are highlighted in red in Figure S6.

*Correction of off-rates to accurately reproduce experiment*. While TPT analysis of our Markov State Model predicts binding on-rates accurately (see Table 1, main text), off-rates are over-estimated by about five orders of magnitude. This is not surprising given the timescales sampling in our simulations. To obtain realistic off-rates for the four-state binding mechanism model, we correct the values of 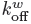 and 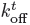 (see Figure 2, main text) according to the experimental off-rate through a scaling factor, γ.

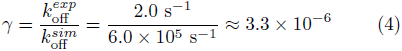

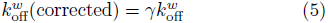

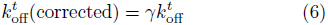

Here, 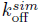 is the off-rate estimated using TPT from the 4-state MSM (see Table 1, main text), and 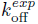 is the experimentally measured off-rate. The original estimates used for correction are from TPT estimation (600-state MSM) as shown in Table S1 and the corrected values are shown in Table S2.

*Correction of p53 folding rates to model residual helicity*. The *helicity* of p53 is defined to be the fraction of population in the helical state:

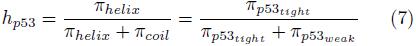

where π_*p*53*tight*_ and π_*p*53*weak*_ are estimated equilibrium populations. The helicity estimated from the MSM model is *h*_*p*53_ = 0.11%.

To model other values of residual helicity in the context of the four-state kinetic model, we must adjust the folding and unfolding rates of unbound p53 TAD peptide, i.e. *k_wt_* and *k_tw_*, respectively. This is an under-constrained problem, as many ratios of rates can lead to the same helix-coil equilibrium constant. Therefore, we turn to a maximum-caliber (MaxCal) approach to infer these rates from the helix/coil population changes, using the relation 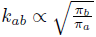 derived in Dixit et al. Given a desired helicity 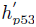, the inferred rates are

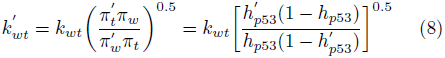

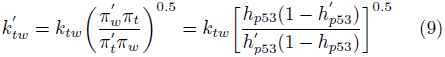

The inferred values are shown in Table S3.

*Derivation of key equations for flux analysis*. The four-state model of Hammes et al. gives expressions for the reactive flux of conformational selection (CS) and induced-fit (IF) pathways in terms of rates and concentrations [p53_Weak_], [p53_tight_], [MDM2 · p53_weak_] and [MDM2]_*f*_, the *free* (unbound) concentration of MDM2. These concentrations can be computed entirely from the rate parameters and the total concentrations [MDM2]_tot_ and [p53]_tot_, as described in Hammes et al. In their original publication, we noticed a typographical error in the expression for [MDM2]_*f*_, and present the correct expression here:

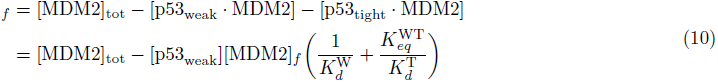

Here, 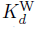 and 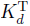 are the dissociation constants of [p53_weak_ · MDM2] and [p53_tight_ · MDM2], respectively; 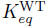 is the equilibrium constant for the reaction p53_weak_ ⇌ p53_tight_.

The concentration of p53_weak_ is calculated using the following equation:

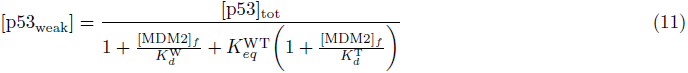

Substituting Eq. 11 into Eq. 10 yields a quadratic equation for [MDM2]_f_:

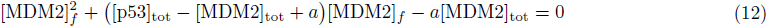

where

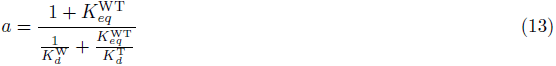

The positive solution of Eq. 12 is the concentration of [MDM2]_*f*_. Once this value is obtained, the concentration of other species can be calculated using the following equations:

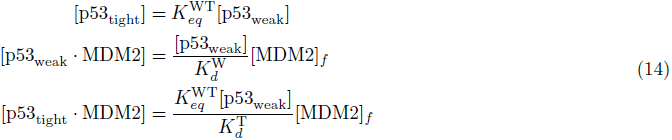

We note that the Hammes et al. model was derived in the context of protein-ligand association, and applied to systems in which the receptor is thought to undergo conformational change that can be either selected or induced by the association of a ligand. In our model, it is p53 that is analogous to the protein receptor, as it undergoes conformational change between a tight-binding helical state and weak-binding coil state, and MDM2 is analogous to the ligand, as it can select or induce helical states of p53 by association.

*Estimation of p53-MDM2 binding affinities at different values of residual helicities*. We calculated the binding affinity ΔG at different values of p53 helicities according to ΔG = *RT*ln *K_d_*. We estimate the disassociation constant *K_d_* based on the four-state model, considering p53_tight_ · MDM2 as the bound state, and the other three states to be unbound states. Thus, the equation used to estimate *K_d_* is

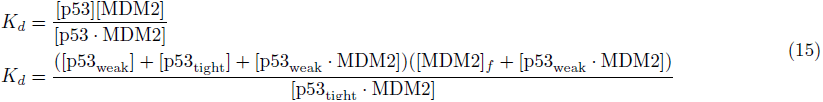

**FIG. S1.**
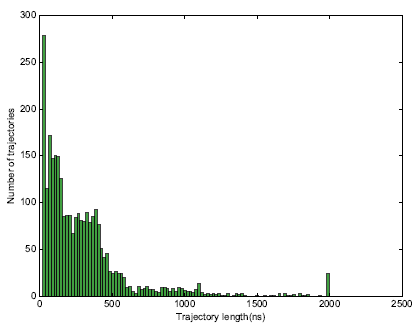
Distribution of simulation trajectory lengths. A total of 2776 trajectories were generated on the Folding@home distributed computing platform, amounting to ~ 831μs of aggregate simulation data.

**FIG. S2.**
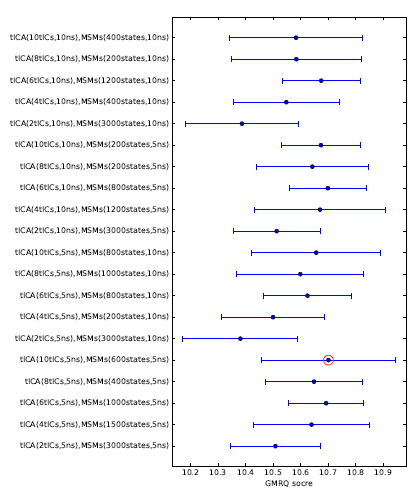
GMRQ scores for MSMs built from various sets of hyper-parameters. Error bars are from 5-fold cross-validation. The hyper-parameters with the best score, marked by a red circle, was chosen for the construction of the final MSM.

**FIG. S3.**
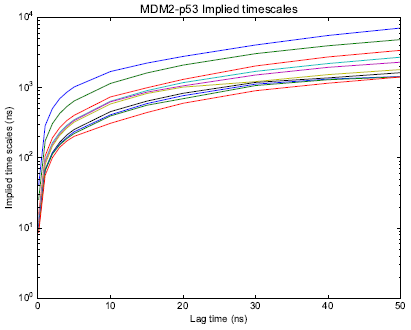
Implied timescales vs MSM lag time. Shown are the ten slowest implied timescales.

**FIG. S4.**
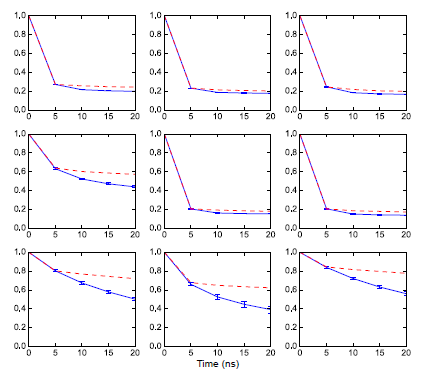
Chapman-Kolmogorov test applied to the nine most populated states in the 600-microstate MSM. Red dashed curves are ⟨*P(i,t* + τ|*i,t*)⟩ for state *i*, calculated from the MD simulations, and blue solid curves are populations [(**T**^(Δ*t*)^)^*n*^´_*i*_]_*i*_ propagated from the MSM transition matrix, where ´_*i*_ is an initial population vector with state *i* containing the entire population, and *n*Δ*t* = τ where τ = 5 ns. Error bars are calculated from five bootstraps.

**FIG. S5.**
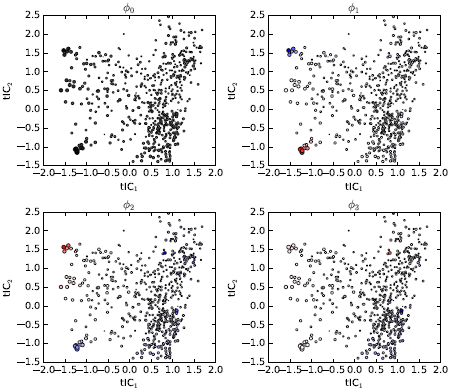
Relaxation eigenmodes of the 600-state MSM. Relaxation dynamics from an initial population vector **p**(0) is given by 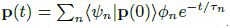, where ψ_*n*_ and ϕ_*n*_ are the left and right eigenvectors of the MSM transition matrix, respectively. Shown are the four slowest relaxation eigenmodes ϕ_*n*_, including the stationary state (i.e. equilibrium populations), ϕ_0_, projected to coordinates tIC_1_ and tIC_2_. The size of each circle is proportional to the equilibrium population, while the color corresponds to the eigenvector structure, with population flux along each eigenmode flowing from blue to red. The slowest relaxation, ϕ_1_ mainly corresponds to binding flux along tIC_1_ while ϕ_2_ corresponds to binding flux along tIC2, both on ~ 1 *μs* timescales (see Figure S3).

**FIG. S6.**
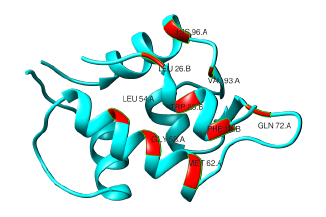
A visualization of residues selected for the computation of p53 distances to the MDM2 binding pocket (see Supporting Methods).

**FIG. S7.**
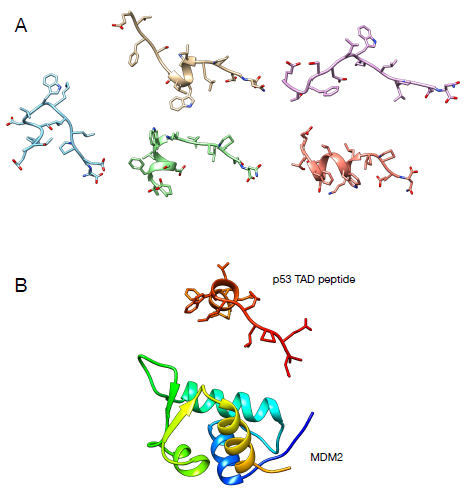
(A) The five initial starting conformations of p53 TAD peptide used in the simulations. These conformations were taken from REMD simulations of p53 TAD peptide in implicit solvent. (B) An example starting configuration of a simulation, with p53 TAD peptide center of mass 12 Å from the binding site.

**TABLE S1.** Estimated rates for flux analysis.

| Parameter | Value (Tmatrix) | Value (TPT) |
| --- | --- | --- |
| $k_{\text{on}}^w$ | $1.8 \times 10^9 \text{ M}^{-1} \text{ s}^{-1}$ | $1.2 \times 10^9 \text{ M}^{-1} \text{ s}^{-1}$ |
| $k_{\text{off}}^w$ | $2.9 \times 10^6 \text{ s}^{-1}$ | $1.8 \times 10^6 \text{ s}^{-1}$ |
| $k_{wt}$ | $1.1 \times 10^4 \text{ s}^{-1}$ | $1.7 \times 10^4 \text{ s}^{-1}$ |
| $k_{tw}$ | $1.0 \times 10^7 \text{ s}^{-1}$ | $9.0 \times 10^5 \text{ s}^{-1}$ |
| $k_{wt}^{\text{MDM2}}$ | $9.6 \times 10^5 \text{ s}^{-1}$ | $1.1 \times 10^6 \text{ s}^{-1}$ |
| $k_{tw}^{\text{MDM2}}$ | $3.9 \times 10^6 \text{ s}^{-1}$ | $4.4 \times 10^6 \text{ s}^{-1}$ |
| $k_{\text{on}}^t$ | $4.0 \times 10^9 \text{ M}^{-1} \text{ s}^{-1}$ | $8.7 \times 10^7 \text{ M}^{-1} \text{ s}^{-1}$ |
| $k_{\text{off}}^t$ | $2.9 \times 10^4 \text{ s}^{-1}$ | $1.8 \times 10^4 \text{ s}^{-1}$ |

**TABLE S2.** Corrected off-rates for the four-state binding mechanism model

| Parameter | Corrected value |
| --- | --- |
| $k_{\text{off}}^w$ | $6.0 \text{ s}^{-1}$ |
| $k_{\text{off}}^t$ | $0.06 \text{ s}^{-1}$ |

**TABLE S3.** Inferred rates for high p53 helicity

| Parameter | Inferred rates |
| --- | --- |
| $k_{wt} \text{ (} h\% = 28\% \text{)}$ | $3.2 \times 10^5 \text{ s}^{-1}$ |
| $k_{tw} \text{ (} h\% = 28\% \text{)}$ | $4.7 \times 10^4 \text{ s}^{-1}$ |
| $k_{wt} \text{ (} h\% = 64\% \text{)}$ | $6.9 \times 10^5 \text{ s}^{-1}$ |
| $k_{tw} \text{ (} h\% = 64\% \text{)}$ | $2.2 \times 10^4 \text{ s}^{-1}$ |

